# *Saccharomyces cerevisiae* displays a stable transcription start site landscape in multiple conditions

**DOI:** 10.1101/393447

**Authors:** Christoph S. Börlin, Nevena Cvetesic, Petter Holland, David Bergenholm, Verena Siewers, Boris Lenhard, Jens Nielsen

## Abstract

One of the fundamental processes that determine cellular fate is regulation of gene transcription. Understanding these regulatory processes is therefore essential for understanding cellular responses to changes in environmental conditions. At the core promoter, the regulatory region containing the transcription start site (TSS), all inputs regulating transcription are integrated. Here, we used Cap Analysis of Gene Expression (CAGE) to analyze the pattern of transcription start sites at four different environmental conditions (limited in ethanol, limited in nitrogen, limited in glucose and limited in glucose under anaerobic conditions) using the Saccharomyces cerevisiae strain CEN.PK113-7D. With this experimental setup we were able to show that the TSS landscape in yeast is stable at different metabolic states of the cell. We also show that the shape index, a characteristic feature of each TSS describing the spatial distribution of transcription initiation events, has a surprisingly strong negative correlation with the measured expression levels. Our analysis supplies a set of high quality TSS annotations useful for metabolic engineering and synthetic biology approaches in the industrially relevant laboratory strain CEN.PK113-7D, and provides novel insights into yeast TSS dynamics and gene regulation.

## INTRODUCTION

Regulation of gene transcription is one of the fundamental processes that determine cellular fate. Transcription of protein encoding genes in eukaryotic cells is governed by the RNA polymerase II in concert with the general transcription initiation factors (GTFs), namely TFIIA, TFIIB, TFIID, TFIIE, TFIIF and TFIIH (reviewed in (1)). These proteins assemble at the core promoter of a gene, which is commonly defined as the minimal region necessary to trigger transcription (2–4). This region encompasses the transcription start site (TSS), defined as the nucleotide position where transcription is initiated (5).

It was previously shown that transcription of a gene in eukaryotic cells is not always initiated from the same nucleotide, but that it can be initiated from a range of positions in the core promoter region, with an individual, sequence-influenced pattern for each gene (3–6). This important finding reshaped the view on transcription initiation showing that there is a higher complexity to this process than previously anticipated.

In addition to the TSS positions being a cornerstone of fundamental knowledge on genome organization, there are numerous applications where an exact mapping of TSS positions is important. One is in the field of synthetic biology, where synthetic promoters are created to obtain a variable range of expression levels. Synthetic promoters are designed by combining core promoters with different upstream regulatory sequences. In order to do this, accurate definition of the promoter regions are needed to place upstream regulatory sequences at the optimal distance to the core promoters. Another application is the modulation of gene expression by CRISPR interference (CRISPRi). An effective strategy for downregulation that has been documented to work for many genes is to target the catalytically inactive Cas9 protein directly to the TSS of the target gene (7).

The most accurate way to map transcription start sites is to selectively sequence intact capped 5’ ends of mRNA. In this study we choose the Cap Analysis of Gene Expression method (CAGE) (8), which was also shown to be the best performing method in a recent comparison of different 5’ end RNA sequencing methods by Adiconis et. al., (9). This method gives a quantitative count of transcription start events with a single base pair resolution, allowing a more detailed interrogation of these events than with traditional RNA sequencing techniques. CAGE can also be used to determine the total expression of a given gene with results showing high correlations with traditional RNA sequencing techniques (10). With this high resolution data it is possible to accurately determine all TSSs of all expressed genes transcribed by RNA Polymerase II and to determine which TSS is the dominant one in a quantitative manner.

Previous work to annotate TSSs has been carried out in different yeast strains using techniques like SMORE-seq (11), or an earlier low-coverage protocol of CAGE (12). These studies used cells grown in shake flasks at only one environmental condition, therefore it was not possible to assess how the TSS landscape changes in response to environmental conditions.

Here we describe the first analysis of the content and dynamics of the yeast promoterome across four different metabolic states. For this, we used an updated CAGE protocol, called non-amplification non-tagging CAGE for Illumina sequencing (nAnT-iCAGE) (13), which is a more unbiased approach compared to the earlier protocol used by Wery et. al. (12) as it omits the use of restriction enzymes to produce short tags and does not include a PCR amplification step of the cDNA. We performed CAGE on the industrially relevant S. cerevisiae laboratory strain CEN.PK113-7D (14), grown in four distinct chemostat conditions at a fixed specific growth rate of 0.1/h. The four chemostat conditions were selected to cover a diverse range of metabolic states, namely: respiratory glucose metabolism using glucose limitation, gluconeogenic respiration using ethanol limitation, aerobic fermentation using nitrogen limitation and fermentative glucose metabolism using anaerobic conditions. With this setup we were able to obtain highly reproducible condition specific data and to assess changes in the TSS landscape in different environmental conditions of the cells.

## MATERIALS AND METHODS

### Gene annotations

To transfer the annotations from the reference genome of S288C (15) to the recently published genome of CEN.PK113-7D (16), first the coding sequences for all verified and uncharacterized ORFs available in the Saccharomyces Genome Database (SGD, www.yeastgenome.org) (17) were obtained using YeastMine, the data API of SGD. Then, using the NCBI software tool Blast+ (18), every obtained sequence was blasted against the CEN.PK genome. Hits covering at least 95% of the sequence length showing at least a 95% sequence identity were retained and transferred if only a single hit existed for that sequence. In case of multiple strong hits, the hit that was found to be on the same chromosome and surrounded by the same neighboring genes as in the reference genome was transferred. In case of large genes where for multiple fragments a hit was found, a manual curation step was performed to check if these fragments could be reassembled into the full gene. Successfully reassembled genes were also transferred, all other hits were discarded. Using this approach, we were able to transfer 99% (5113 out of 5159) of the verified ORF annotations and 96% (727 out of 756) of all uncharacterized ORF annotations. The gene YCL018W, which was found to be duplicated in the originally published sequence, was also found to be duplicated using our approach (16). In addition to that, the genes YHR055C and YHR054C were also found to be duplicated. The complete set of updated annotations can be found in the supplementary table S1.

### Chemostat cultures and RNA extraction

The S. cerevisiae strain CEN.PK113-7D (14) was pre-cultured in a batch culture in 100 ml of minimal medium with 2% glucose (See Table 1 for media composition and recipe) at 30°C and 200 rpm in 250 ml shake flasks for 24 hours. The pre-culture was then transferred to the chemostats in triplicates to an initial OD600 above 3. For each of the four conditions a single pre-culture was used. Four different media compositions with a limitation in a different nutrient were employed in the chemostat runs (See Table 1 for media composition and recipe). The medium volume in the chemostat runs was 40 ml and the temperature was set to 30°C. One hour after the transfer the pumps where started with the dilution rate fixed to 0.1/h. Dissolved oxygen was kept above 30% of air saturation. For the anaerobic condition, the fermenter was flushed with nitrogen gas. The cells were grown for 4 days to achieve stable cell numbers in the culture. An amount of cells corresponding to a total OD600 of 10 were collected, pelleted and snap frozen in liquid nitrogen for both RNAseq analysis and the CAGE experiment.

**Table 1.** Synthetic media composition for the four different chemostat conditions and the batch preculture. The amounts indicated are for one liter of media.

|  | Batch Pre-Culture | Ethanol limited | Glucose limited | Nitrogen limited | Anaerobic glucose limited |
| --- | --- | --- | --- | --- | --- |
| KH<sub>2</sub>PO<sub>4</sub> | 14.4 g | 14.4 g | 14.4 g | 14.4 g | 14.4 g |
| MgSO<sub>4</sub> | 0.5 g | 0.5 g | 0.5 g | 0.5 g | 0.5 g |
| (NH<sub>4</sub>)<sub>2</sub>SO<sub>4</sub> | 7.5 g | 5 g | 5 g | 1 g | 5 g |
| K<sub>2</sub>SO<sub>4</sub> | - | - | - | 5.3 g | - |
| Glucose 40% | 50 ml | - | 18.75 ml | 150 ml | 25 ml |
| Ethanol 96% | - | 6.7 ml | - | - | - |
| Trace metals stock solution | 1 ml | 1 ml | 1 ml | 1 ml | 1 ml |
| Vitamin stock solution | 1 ml | 1 ml | 1 ml | 1 ml | 1 ml |
| Ergosterol in Tween 80 (2.6 g/L) | - | - | - | - | 4 ml |
| Antifoam 204 | - | 7 drops | 7 drops | 7 drops | 9 drops |

Trace metal stock solution components (per liter of stock solution): 15.0 g EDTA-Na<sub>2</sub>, 4.5 g CaCl<sub>2</sub>·2H<sub>2</sub>O, 4.5 g ZnSO<sub>4</sub>·7H<sub>2</sub>O, 3 g FeSO<sub>4</sub>·7H<sub>2</sub>O, 1g H<sub>3</sub>BO<sub>3</sub>, 0.84 g MnCl<sub>2</sub>·2H<sub>2</sub>O, 0.4 g Na<sub>2</sub>MoO<sub>4</sub>·2H<sub>2</sub>O, 0.3 g CuSO<sub>4</sub>·5H<sub>2</sub>O, 0.3 g CoCl<sub>2</sub>·6H<sub>2</sub>O and 0.1 g KI. Vitamin stock solution components (per liter of stock solution): 25 g myo-inositol, 1 g nicotinic acid, 1 g calcium pantothenate, 1 g pyridoxine HCl, 1 g thiamine HCl, 0.2 g 4-aminobenzoic acid and 0.05 g biotin. The pH of the media was adjusted to 6.0-6.5 using KOH pellets, resulting in a final pH around 5.5 in all four chemostat cultures.

Cells were mechanically disrupted using a FastPrep®-24 from MPbio (Santa Ana, California, USA) in combination with the lysing matrix tubes type C from MPbio. The FastPrep was run 3 times for 20 seconds with 4.0m/s settings and a 5 minute break in between each run. RNA was subsequently extracted using the RNeasy^®^ Mini Kit from QIAGEN (Hilden, Germany). RNA quality was assessed using ThermoFischer NanoDrop (Waltham, Massachusetts, USA) and Agilent2100 Bioanalyzer (Santa Clara, California, USA) to ensure high quality RNA.

## RNA sequencing

All three biological replicates for each condition were sequenced using the NextSeq500 System from Illumina (San Diego, California, USA) at the Novo Nordisk Foundation Center for Biosustainability, Technical University of Denmark, with paired-end reads of 75 bp length. Library preparation was done using the Illumina TrueSeq stranded total RNA HT kit following the manufacturer’s instructions. Obtained reads were mapped to the CEN.PK113-7D genome using bowtie2 (19). Mapped reads were filtered using a quality threshold of 20 and converted to .bam files using samtools (20). FeatureCounts was used to obtain expression values for each gene (21), which were subsequently converted into TPM values.

### CAGE

For the CAGE experiment the non-amplification non-tagging CAGE protocol for Illumina sequencing (nAnT-iCAGE) as previously published by Murata et. al. (13), was used on two biological replicates of each condition, starting with 5 micrograms of extracted total RNA. The 8 barcoded samples were pooled together and sequenced using the Illumina HiSeq 2500 at Genomics Core Facility (MRC, London Institute of Medical Sciences). Between 2.6 to 22.1 million reads per sample were obtained, with an average of 9 million, showing a very high coverage of the yeast transcriptome. Sequencing reads were mapped to the CEN.PK113-7D genome using bowtie2 (19). Mapped reads were filtered using a quality threshold of 20 and converted to .bam files using samtools (20). 29% of all reads mapped to a 7.2 kb region with ribosomal repeats on chromosome 12, which was excluded from further analysis, leaving an average mapped read count of 5.4 million reads per replicate. An overview of the sequencing read numbers is shown in Supplementary table S2.

CAGE data were analyzed using the R/Bioconductor package CAGEr (22, 23). The .bam files were imported into R and the biological replicates were merged together. Default CAGEr correction of the first G nucleotide was used. The data were then normalized for library size using the “powerLaw” method (24) with a fit range from 5 to 10000 and an alpha value of 1.10. The CAGE tags were clustered together using the “clusterCTSS” function of CAGEr with the “distclu” setting, a maximum distance of 20 and a TPM threshold of 1. These clusters were then aggregated across the conditions to obtain a set of consensus clusters using the “aggregateTagClusters” function with a TPM threshold of 3 and a maximum distance of 100. For each consensus cluster the expression level in every condition was calculated as TPM and the dominant TSS position was calculated based on the normalized tag count per base. Additionally, the shape index of each cluster was calculated by the formula described by Hoskins et. al. (25): 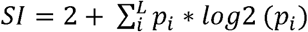. A graphical example for this with two artificial promoters can be found in Supplementary Figure S3.

*p_i_* = proportion of counts at position i in the cluster

*L* = position with at least 1 tag

### Annotation of CAGE clusters to genes

The obtained CAGE consensus clusters were assigned to the gene annotations using the following set of rules: The consensus cluster must be on the same strand as the gene annotation, the cluster is not more than 1 kb away from the start of the gene annotation and if the cluster is upstream of the gene annotation, RNAseq reads must be present covering the region between the cluster and the gene annotation.

### Expression based clustering of genes

For creating the four gene clusters based on the expression profiles we normalized the expression across the four conditions and a gene was assigned to one of the four clusters if it meets the following requirements: Cluster “Always”: The expression level in each condition must account for 23 to 27% of the total observed expression. Cluster “Glu+Eth”: At least 83.3% of the observed total expression must come from the two respiratory conditions (respiratory glucose metabolism using glucose limitation and gluconeogenic respiration using ethanol limitation). Cluster “Nit”: At least 75% of the observed total expression must come from aerobic fermentation using nitrogen limitation. Cluster “Ana”: At least 75% of the observed total expression must come from fermentative glucose metabolism using anaerobic conditions.

## RESULTS AND DISCUSSION

### CAGE data are highly reproducible and reveal promoters and TSS for 88% of all annotated genes

In order to gain insights into the promoter structure of the yeast strain CEN.PK113-7D and to obtain accurate positions of the transcription start sites (TSS) we performed a cap analysis gene expression (CAGE) experiment on yeast grown in four different chemostat conditions. The conditions were: respiratory glucose metabolism using glucose limitation (Glu), gluconeogenic respiration using ethanol limitation (Eth), aerobic fermentation using nitrogen limitation (Nit) and fermentative glucose metabolism using anaerobic conditions (Ana).

Using the R/Bioconductor package CAGEr (22), for each of the four conditions we assembled the single position read tags into clusters and then merged overlapping clusters together to form a consensus cluster, combining information from all four conditions. This resulted in a total of 6565 consensus clusters which were then assigned to the gene annotations. For 5247 clusters a matching gene annotation could be found, of which 4975 genes were assigned a single cluster and 133 genes were assigned multiple clusters (ranging from 2 to 4 clusters per gene, a total of 272 clusters). This means that from a total of 5843 gene annotations in the genome, we could assign 5108 (88%) to at least one cluster. The complete set of results for each individual cluster can be found in supplementary table S4. A representative display of the CAGE data, in the Integrative Genomics Viewer (IGV) (26, 27), is shown in Figure 1A-D. The four genes were selected to showcase different distributions of CAGE reads for condition independent as well as condition specific genes. Figure 1A and 1B show genes that are expressed in all four conditions, while Figure 1C and 1D show genes that are only expressed in some of the conditions. In addition to that, Figure 1A and 1C show genes with a broad TSS distribution, while Figure 1B and 1D show genes with a peaked TSS distribution, showing that a broad or peaked TSS distribution is not unique for condition independent or specific genes.

**Figure 1.**
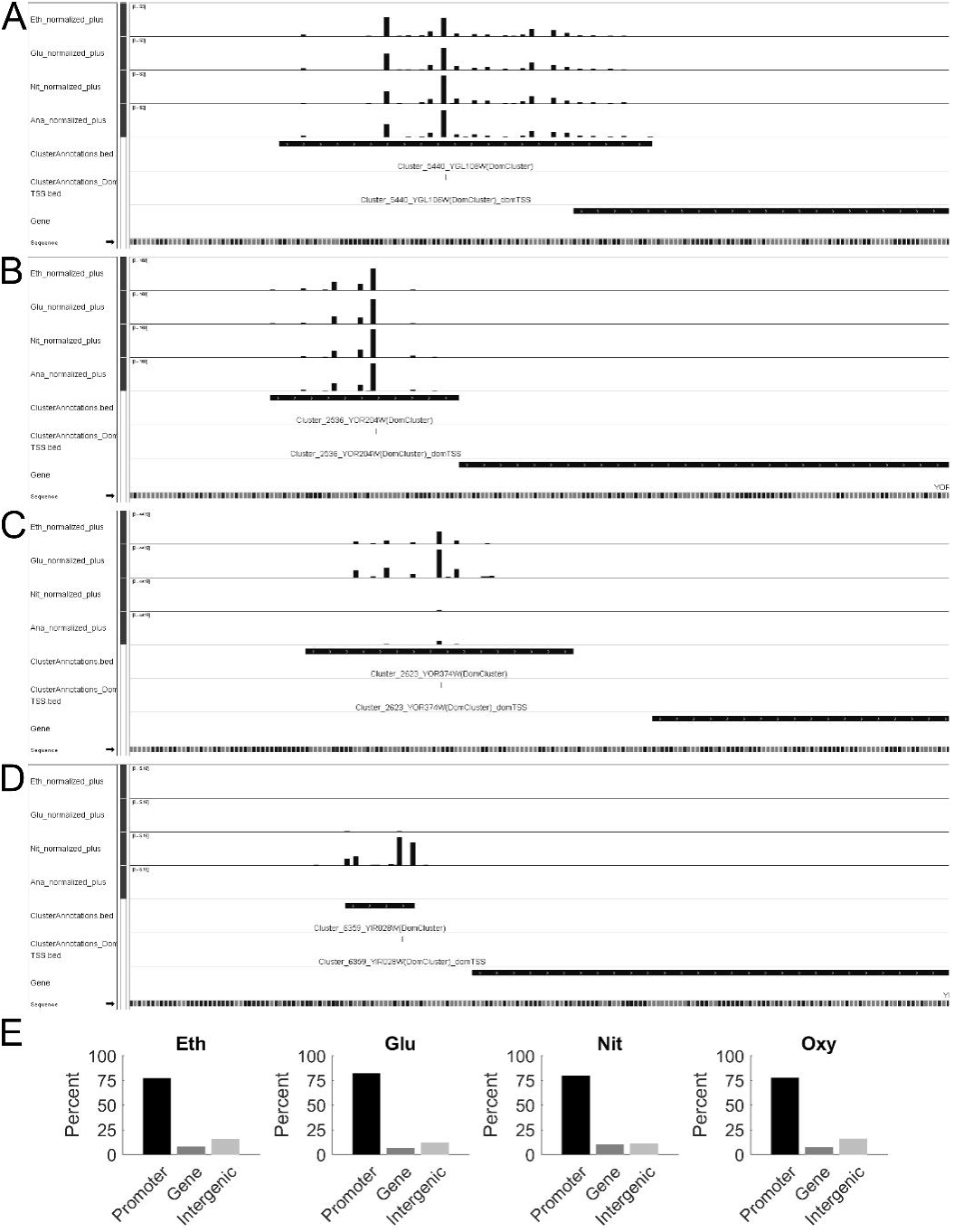
Overview of the obtained CAGE data. A Screenshot from IGV showing the broad CAGE read distribution for the constitutively expressed gene YGL106W (MLC1). B Screenshot from IGV showing the peaked CAGE read distribution for the constitutively expressed gene YOR204W (DED1). C Screenshot from IGV showing the broad CAGE read distribution for a gene mainly expressed in the respiratory conditions YOR374W (ALD4). D Screenshot from IGV showing the peaked CAGE read distribution for a gene exclusively expressed in aerobic fermentation YIR028W (DAL4). E Intersection of mapped CAGE reads with gene annotations, the promoter region was defined as the 500bp upstream of the start of the coding region and which was therefore not considered to be part of the intergenic region. (Eth = gluconeogenic respiration using ethanol limitation, Glu = respiratory glucose metabolism using glucose limitation, Nit = aerobic fermentation using nitrogen limitation, Ana = fermentative glucose metabolism using anaerobic conditions).

To assess the quality of our obtained CAGE data, we first analyzed the location of the sequencing reads in relation to the annotated genes (Figure 1E) and found that the majority of all reads (77% to 82%) map to the promoter region of annotated genes, which was defined as the 500 bp region upstream of the start of the coding sequence. As the TSS of a gene is expected to be upstream of the coding sequence, this also indicates that we obtained high quality CAGE data.

179 clusters were annotated as possible antisense transcription events to a total of 169 genes, as they were located at the 3’ end of the gene on the opposite strand. It has been shown before that wide-spread anti-sense transcription occurs in yeast (28). Yassour et al. reported how much of every gene was covered by anti-sense transcription, and in their study 1523 genes had at least 10% of their sequence covered by anti-sense reads. Comparing these 1523 genes with our list of 169 genes with possible anti-sense initiation we find a high degree of overlap of 75% (122 out of 163 genes that are in both lists). As this finding is in line with the already known wide-spread anti-sense transcription, we focused on the sense transcription events. From the total of 6565 consensus CAGE clusters, 1115 clusters could not be assigned to any gene.

To further check the quality of the obtained CAGE data, the individual samples were clustered together using hierarchical clustering based on their genome-wide expression profile at each base pair. For each condition the replicates cluster together (Figure 2A), showing the high reproducibility of the data. This clustering also shows that the two respiratory conditions (respiratory glucose metabolism using glucose limitation, gluconeogenic respiration using ethanol limitation) are more similar to each other than to the two fermentative conditions, as one would expect. In addition, the correlations between the biological replicates were calculated and with a minimum Pearson correlation coefficient of 0.9 the replicates are in high agreement with each other (Figure 2A).

**Figure 2.**
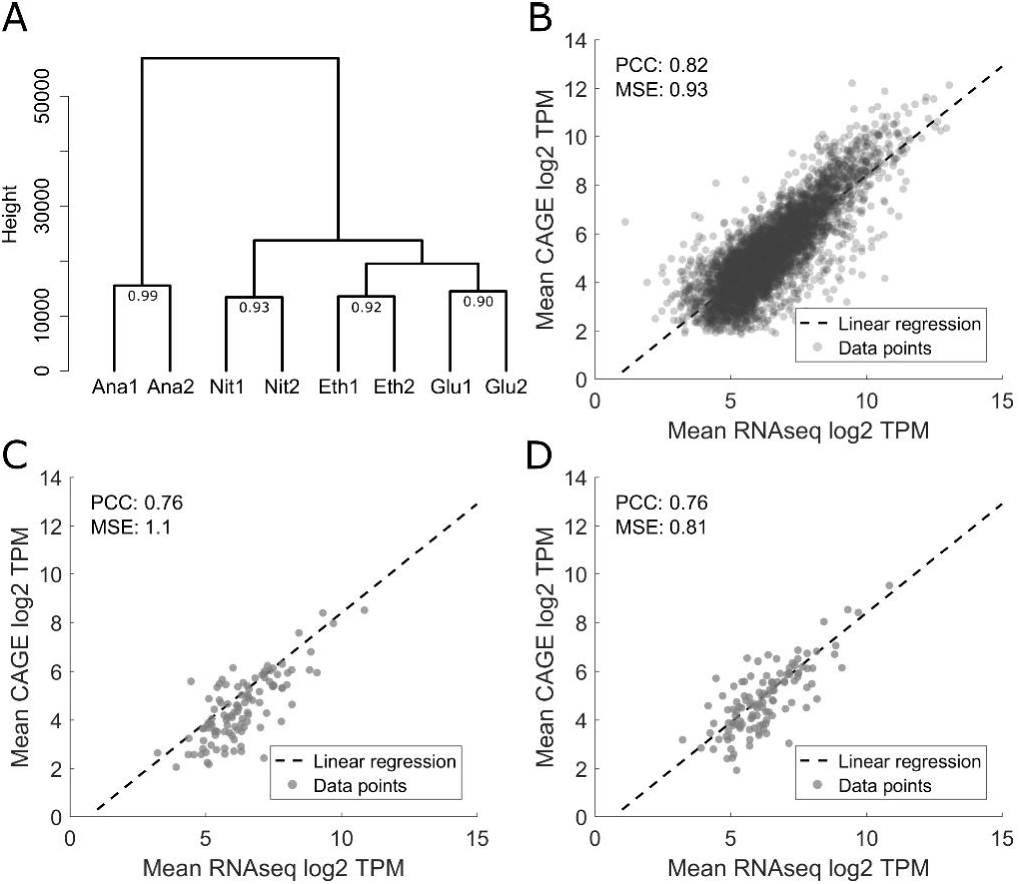
Quality control of CAGE expression levels by comparison with RNAseq expression levels A Hierarchical clustering of the individual CAGE sequencing experiments based on normalized TSS tag values. The number at the last branch points denotes the Pearson correlation between the replicates B Comparison between average expression values across the conditions obtained through RNAseq and CAGE, both in log2 TPM values. Only genes with a single CAGE cluster were compared, showing a high correlation with a Pearson correlation coefficient of 0.82. Red lines show the linear regression fitted to this data, with a mean squared error (MSE) of 0.93. C Comparison of RNAseq and CAGE expression data for genes with more than one CAGE cluster, where only the strongest CAGE cluster was used for calculating the TPM value, resulting in an MSE of 1.1 and a PCC of 0.76. D Comparison of RNAseq and CAGE expression data for genes with more than one CAGE cluster. All CAGE clusters where added up to calculate the TPM value, resulting in an MSE of 0.81 and a PCC of 0.76.

Subsequently the expression values per gene promoter region obtained from CAGE was compared with gene expression values obtained from a control RNAseq experiment in the same chemostat conditions (Figure 2B-D). Comparing genes with a single annotated cluster shows a high correlation of the two expression values, with a Pearson correlation coefficient of 0.82 (Figure 2B). In order to obtain a conversion formula between RNAseq and CAGE values, a linear regression was fitted to the data (Figure 2B, dotted line). This enabled us to then compare the expression values of genes with multiple annotated clusters in two different ways: considering only the strongest cluster of each gene (Figure 2C) or summing up the expression values from all CAGE clusters per gene (Figure 2D). As expected, taking all clusters into consideration leads to a better fit to the previously established conversion formula. However, the correlation between the CAGE and RNAseq expression values for genes with multiple clusters is lower than for single cluster genes. One reason could be that for genes with multiple clusters the calculation of RNAseq TPM values is not correct, because it assumes a uniform gene length, while in reality there are two or more differently sized transcripts present in the cells.

### The yeast TSS landscape shows stability across metabolic conditions in a variety of characteristics

One possible analysis of CAGE clusters is to look at the cluster width or to look at the interquantile cluster width for the quantiles 0.1 to 0.9, as established by Haberle et. al. (22). The interquantile cluster width is more robust to noise and therefore this approach was chosen for subsequent analysis. For each gene, an average cluster width across the four conditions was calculated. The distribution of the average widths (Figure 3A) shows a unimodal distribution with an average width of 31 bp. The interquantile cluster width is also stable across the four conditions, as shown in a pairwise comparison between conditions, with a minimum Pearson correlation coefficient of 0.84 (Figure 3B).

**Figure 3.**
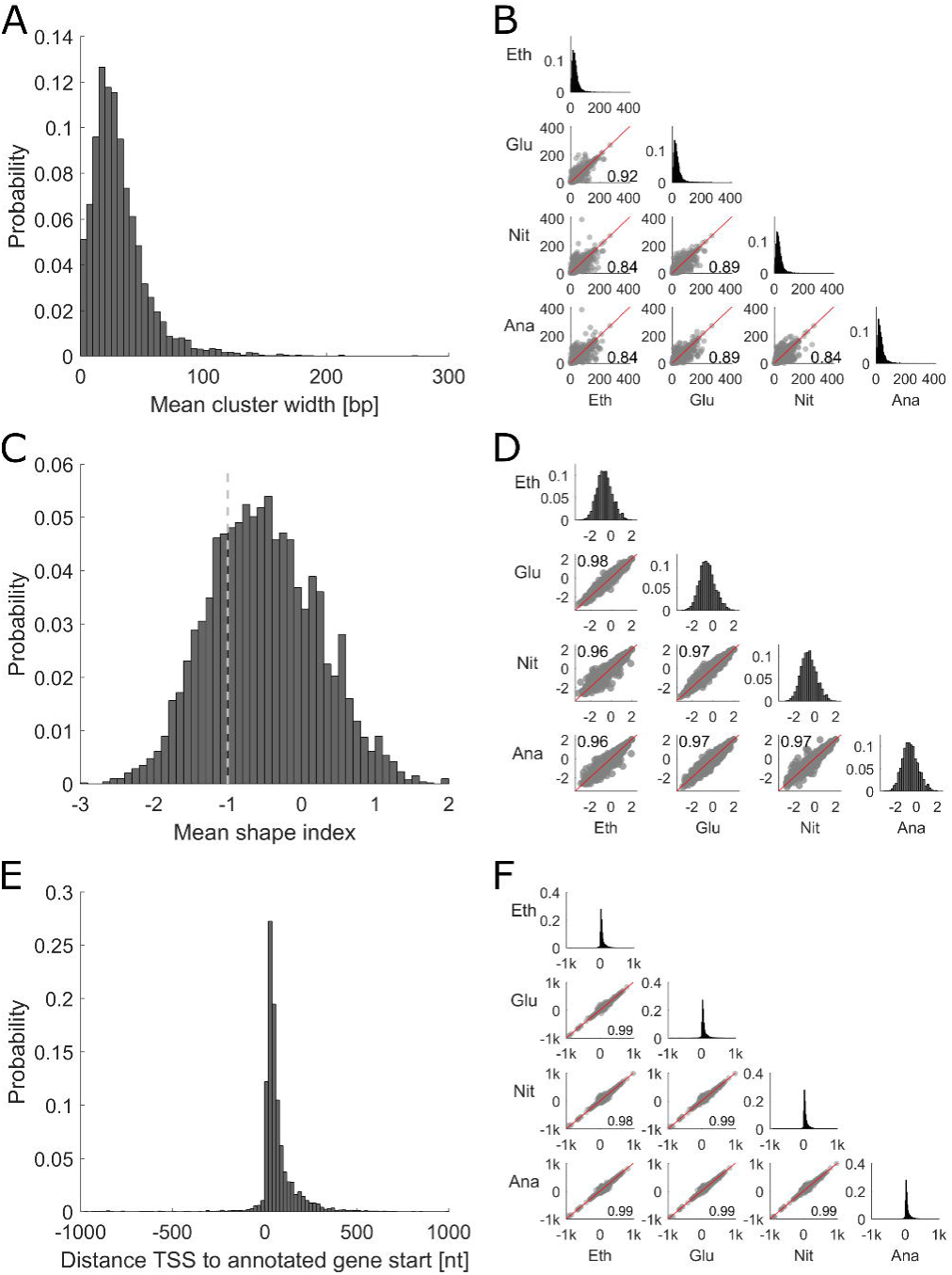
Overview of TSS cluster characteristics and their stability across conditions. A Histogram showing distribution of mean interquantile cluster width across all conditions. B Comparison of the promoter width in different conditions. Middle axis showing the distribution in each condition and the lower half displaying the pairwise comparison of each condition together with the Pearson correlation coefficient. C Histogram showing the distribution of distances between the global TSS across conditions and the assigned genes. D Comparison of the distance between the condition specific TSS and the assigned gene in different conditions. Middle axis showing the distribution in each condition and the lower half displaying the pairwise comparison. E Histogram showing the mean shape index of each cluster. The dashed line at -1 denotes the border which separates clusters classified as peaked (shape index >-1) and clusters classified as broad (shape index <= -1). F Comparison of the shape index in different conditions. Middle axis showing the distribution in each condition and the lower half displaying the pairwise comparison.

It has been shown that the cluster width is not always sufficient to classify clusters as either peaked or broad. Clusters that are very wide but where the dominant positions contribute the majority of reads exist, as well as narrow clusters with multiple near equally strong positions. To overcome this issue, Hoskins et. al. established the shape index as a more informative tool for this classification (25). The formula used to calculate the shape index can be found in the method section and an example classification of two artificial clusters is shown in the supplementary Figure S3. In short, a gene showing a peaked TSS distribution will obtain a shape index higher than -1, while a gene showing a broad TSS distribution will obtain a value lower than or equal to -1. We classified the peaks detected by our analysis by the average shape index across the conditions and we find that the majority of clusters are classified as peaked, i.e. have a shape index higher than -1 (Figure 3C). The shape index is also very stable between the conditions, with a minimal pairwise Pearson correlation coefficient of 0.96 (Figure 3D).

Calculating the 5’ UTR length (Figure 3E) showed that most clusters are quite close to the start of the coding sequence, with 70% of them being less than 75 bp away, which is in line with previously published average 5’ UTR lengths in yeast (11). This 5’ UTR length is again very stable across the different conditions with a minimal Pearson correlation coefficient of 0.99 between two individual conditions (Figure 3F). We further compared the published TSS dataset from Parky et. al. (11), obtained using the yeast strain BY4741 in YPD, with our TSS annotations for CEN.PK113-7D. For the 4872 genes that are present in both data sets we calculated the 5’ UTR lengths (using sacCer3 annotations for Parky’s dataset and our CEN.PK113-7D annotations for our dataset) and compared them. The 5’ UTR lengths are in high agreement with each other, with an average difference of less than 9 bp. Both datasets have around 250 TSS annotations for genes not found in the other dataset, these differences are most likely due to different expression profiles caused by different media and growth conditions (YPD in shake flasks vs synthetic minimal media in chemostats) or strain differences (BY4741 vs CEN.PK113-7D). The high agreement between the two datasets highlights the quality of our TSS annotations for the industrially relevant strain CEN.PK113-7D.

### Gene clustering by condition specific expression shows no distinct promoter characteristics

To further test the stability of the yeast transcriptional landscape in different conditions we clustered genes together based on their expression levels in different conditions. For this we normalized the genes expression levels across the conditions and created the following four gene clusters: 1: Genes that are expressed under all four conditions (labeled as “always”); 2: Genes that are mostly active in the two respiratory conditions (respiratory glucose metabolism using glucose limitation and gluconeogenic respiration using ethanol limitation, labeled “Glu+Eth”); 3: Genes that are mainly active in aerobic fermentation using nitrogen limitation (labeled “Nit”) and 4: Genes that are mainly active in fermentative glucose metabolism using anaerobic conditions (labeled “Ana”). For each of these four groups we analyzed the expression levels (Figure 4A), the interquantile widths (Figure 4B), the shape indices (Figure 4C) and the 5’ UTR length (Figure 4D). The overall picture shows that there are no remarked differences in these characteristics between the four groups. The average gene expression levels are quite similar, with the genes expressed in all four conditions showing a slightly narrower distribution than the condition specific genes, a trend that can also be seen in the distribution of shape indices. These differences however are not very strong.

**Figure 4.**
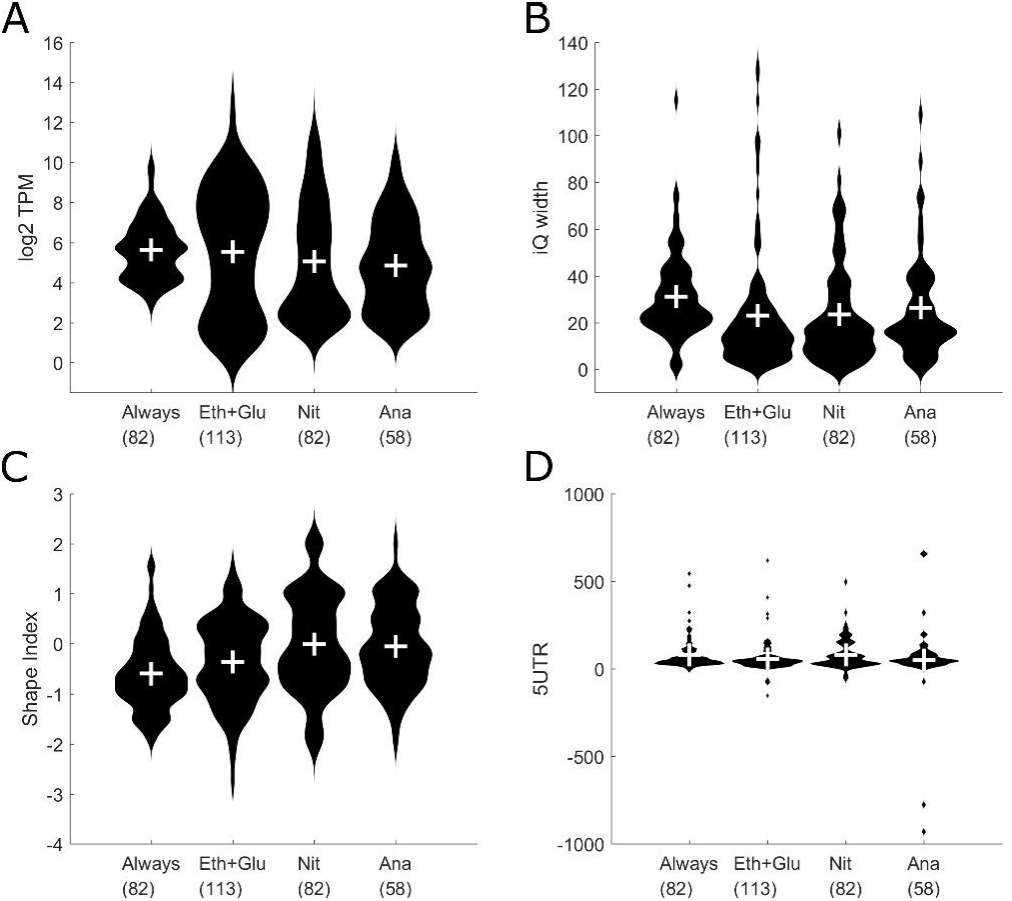
Comparison of condition based gene clusters (Always = Genes that are expressed in all four conditions, Eth+Glu = Genes active in both respiratory conditions, Nit = Genes active under aerobic fermentation, Ana = Genes active in fermentative glucose metabolism). The number under the gene cluster denotes the number of genes in that cluster. A Violin plot showing distribution of gene expression levels for each gene cluster. B Violin plot showing distribution of the interquantile promoter width for each gene cluster. C Violin plot showing distribution of the shape index for each gene cluster. D Violin plot showing distribution of the 5’ UTR lengths.

### Cluster shape shows a high correlation to promoter expression levels

In higher organisms like Drosophila melanogaster, there is a remarkable relation between the shape index of a TSS cluster and the gene expression level during different developmental phases (25). Genes with a broad TSS cluster show a stable expression level throughout embryonic development, while genes with a peaked cluster show a transcription pattern that varies in time and space (25). To see if this relationship between shape index and gene expression variability also holds for yeast, we averaged the shape index of each cluster across the four conditions and then selected the 100 genes with the lowest shape index, i.e. the genes with the broadest clusters, and the 100 genes with the highest shape index, i.e. the genes with the most peaked clusters. For these selected genes we then compared the expression values in each individual condition (Figure 5A-B). No significant differences in expression levels were observed when comparing the four different conditions. However, there was a marked difference in the overall expression levels between genes with a broad cluster and genes with a peaked cluster (comparing overall TPM levels in Figure 5A with 5B). Following this observation, the correlation between the mean shape index across the conditions and the mean expression levels was analyzed, as shown in Figure 5C. A striking anticorrelation with a Pearson correlation coefficient of -0.45 was observed, indicating that peaked clusters (clusters with a high shape index) in yeast are associated with lower expression levels. To check if that strong correlation was unique to the shape index, we also calculated the correlation between the mean interquantile promoter width with gene expression levels (Figure 5D) and we observe no correlation. This indicates that the strong correlation observed between the shape index and gene expression levels is a unique feature of the shape index.

**Figure 5.**
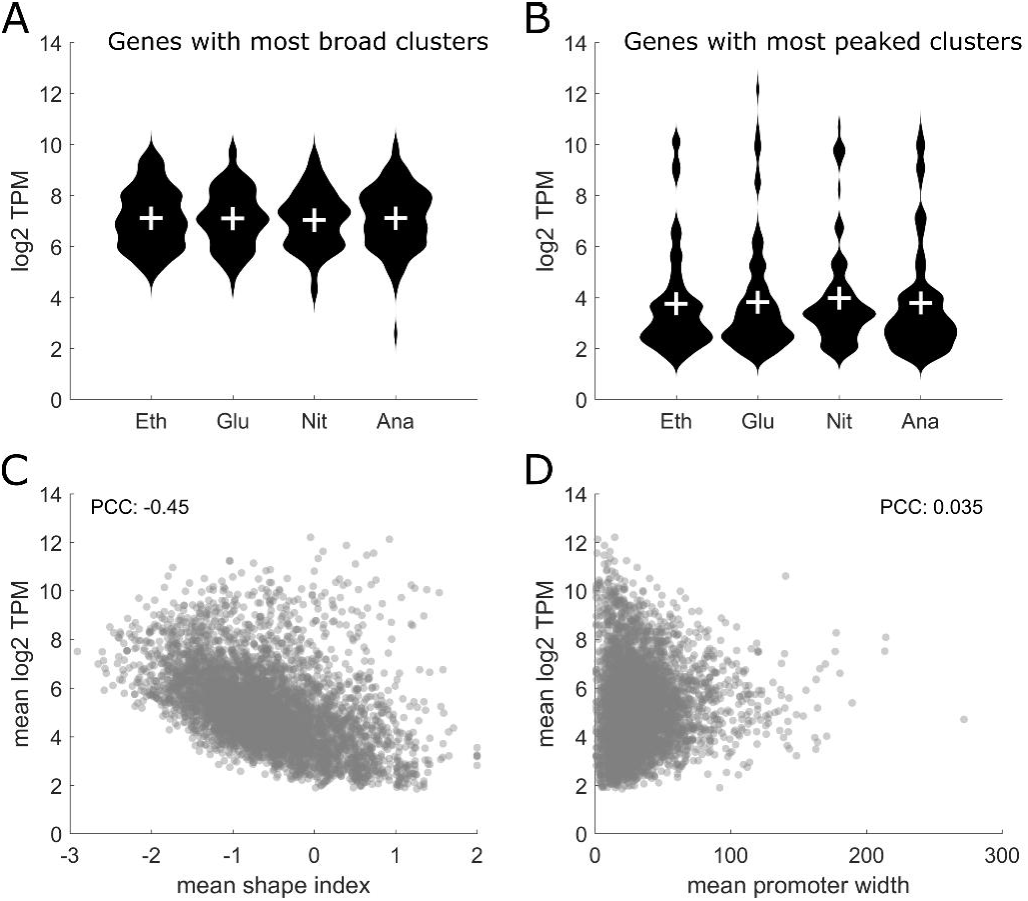
Detailed analysis of the Shape Index. A The 100 genes with the broadest clusters across all conditions were selected and their expression values in each condition are shown. B The 100 genes with the most peaked clusters across all conditions were selected and their expression values in each condition are shown. C Correlation of the mean shape index and the mean CAGE expression levels showing an anticorrelation with a Pearson correlation coefficient of -0.45. D Correlation of the mean promoter width with mean CAGE expression levels showing no correlation.

### Data availability and usage

To enable the easy usage of our data, we created custom data tracks and sessions for the Integrated Genomics Viewer (IGV, (26, 27)), which can be found in the supplementary data. After downloading the IGV from http://software.broadinstitute.org/software/igv/home, first the CEN.PK113-7D genome file (“CEN.PK113-7D.genome” part of the zipped supplementary file S5) has to be loaded via “Genomes” -> “Load Genome from File…” menu in IGV. After that, it is possible to load the session file for either the raw CAGE reads (“IGV_session_RawData.xml”, part of the zipped supplementary file S6) or the normalized CAGE reads (“IGV_session_NormData.xml”, part of the zipped supplementary file S7) using the “File” -> “Open Session…” menu.

After loading the session, a screen similar to the one shown in Figure 1A-D will be visible. For each condition there are two tracks, one for reads on the plus strand and one for reads on the minus strand of the genome (labeled “_plus” and “_minus” respectively). In addition there are three different annotation tracks. The first one, labeled “ClusterAnnotations.bed”, will show each cluster with the full width, while the second one, labeled “ClusterAnnotationsDomTSS.bed”, will only show the position of the strongest TSS in each cluster. Both of these tracks include information about the cluster ID, and to which gene the cluster is annotated to (if any). For each gene, the strongest cluster is labeled as “(DomCluster)”. A third annotation track called “Gene” displays the blast based gene annotations for the CEN.PK113-7D genome.

## Conclusion

In this study, we present a high quality CAGE dataset in four distinct chemostat conditions, to accurately annotate the TSS of each gene. This resource will be valuable to the community as accurate TSS annotations, based on the dominant TSS position, are valuable for promoter engineering and implementation of CRISPRi approaches.

Analysis of the yeast promoterome in the different conditions shows a remarkable level of stability in terms of promoter characteristics like promoter width and shape index. This is in contrast to higher organisms where strong changes can occur, especially during embryonal development stages (29), and suggests that the basic regulatory events governing gene expression in yeast are quite distinct from other eukaryal cells.

## ACCESSION NUMBERS

The complete CAGE sequencing data and results can be found under the ArrayExpress accession code E-MTAB-6650 after publication.

The complete RNA sequencing data and results can be found under the ArrayExpress accession code E-MTAB-6722 after publication

## SUPPLEMENTARY DATA

Supplementary Data will be available online after publication.

